# Spectroscopic study on binding of cationic Pheophorbide-*a* to antiparallel quadruplex Tel22

**DOI:** 10.1101/462291

**Authors:** Olga Ryazanova, Victor Zozulya, Igor Voloshin, Alexander Glamazda, Igor Dubey, Larysa Dubey, Victor Karachevtsev

## Abstract

Binding of water-soluble cationic Pheophorbide-*a* derivative (CatPheo-*a*) to Na^+^–stabilized antiparallel quadruplex formed by 22-mer oligonucleotide d[AG_3_(T_2_AG_3_)_3_], a fragment of human telomeric DNA (Tel22, PDB ID: 143D), has been examined using experimental techniques of absorption and polarized fluorescent spectroscopy as well as absorption melting. The binding affinity of CatPheo-*a* to Tel22 was studied in titration experiments registering the dependence of the dye fluorescence intensity and polarization degree on molar phosphate-to-dye ratio (*P/D*). CatPheo-*a* was found to bind effectively to the quadruplex, and two competitive binding modes were detected. The first one predominates at the dye excess and results in the fluorescence quenching, whereas the second one is preferential at the biopolymer excess and results in the enhancement of pheophorbide emission. The effect of CatPheo-*a* on thermodynamic parameters of Tel22 quadruplex unfolding was estimated using a two-state model. It was found that CatPheo-*a* destabilizes the quadruplex structure of Tel22 slightly decreasing its 4→1 transition midpoint temperature, gives destabilizing increment into Gibbs standard free energy and 2-fold decrease in the equilibrium quadruplex folding constant at 37°C. In ethanol CatPheo-*a* exhibits 15% higher efficiency of singlet oxygen generation as compared to the parent Pheo-*a* compound that makes it a promising photosensitizer for photodynamic therapy.

## 1. Introduction

Porphyrins represent a large class of macrocyclic compounds possessing by unique photophysical properties and having the great potential for application in biology, medicine and nanotechnology. One of them, Pheophorbide-*a* (Pheo-*a*), is a well-known chlorophyll metabolite characterized by the chlorine-type visible absorption spectrum [1,2] with high extinction coefficient in the red region where the transparency of tissues to light increases considerably. This dye selectively accumulates in the tumor cells that determine its widespread using as a photosensitizer for PDT of tumors [3–6]. One of the mechanisms of the Pheo-*a* action on tumor tissues is based on the ability of the dye to produce singlet oxygen in cells under red light irradiation [7]. However, natural Pheo-*a* has poor water solubility. It contains a carboxylic group, which anionic character complicates binding of the dye to negatively charged nucleic acids. To correct this disadvantage a novel cationic derivative of this dye containing a trimethylammonium group (CatPheo-*a*, figure 1) was synthesized by us [8]. We supposed that transformation of Pheo-*a* into cationic compound would improve its DNA binding ability.

**Figure 1.**
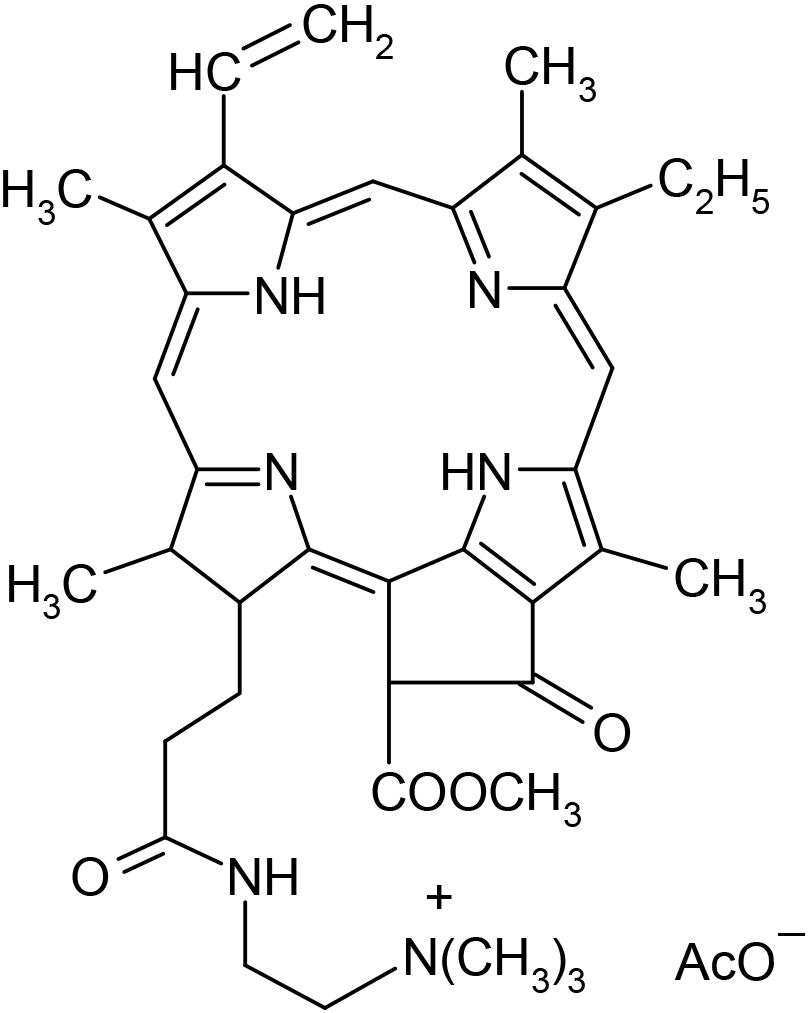
Molecular structure of CatPheo-*a*.

Spectroscopic properties of CatPheo-*a*, as well as the ways of its binding to model synthetic biopolymers of different secondary structures (single-strand inorganic polyphosphate, double-stranded poly(A)⋅poly(U), poly(G)⋅poly(C) polynucleotides and four-stranded poly(G) polynucleotides) were studied earlier at near-physiological ionic conditions using various spectroscopic techniques [8,9]. It was shown that CatPheo-*a* binds to all these polymers, but especially effective binding was observed in the case of quadruplex poly(G), evidently due to the structural features of the dye core which is appropriate for interaction with the G-tetrads through π – π stacking interactions. Two types of CatPheo-*a* complex with long four-stranded poly(G) were revealed depending on the relative phosphate–to–dye concentration ratio, *P/D* [9]. The first one predominating at high dye content (low *P/D*) and characterized by weakening of the dye emission was identified as external binding of the dye to phosphate backbone with self-stacking. The second binding mode prevalent at high *P/D* (presumably, intercalative binding) was accompanied by the enhancement of CatPheo-*a* fluorescence. High affinity revealed for CatPheo-*a* binding to guanine-rich nucleic acids gives the ground to assume the improved antitumor properties of the dye.

G-quadruplexes (G4) are specific four-stranded nucleic acid structures containing π-π stacked guanine tetrads, normally stabilized by metal cations located in between the tetrads to coordinate the guanine bases. They are considered as key regulatory elements in genome regions of biological significance, in particular, in human telomeres and promoter regions of some proto-oncogenes [10-13]. They are known as promising targets for anticancer therapy, since G-quadruplex binding ligands, including many porphyrin derivatives, are able to inhibit the telomerase, a tumor-associated enzyme responsible for telomere elongation [14–16]. We have recently found that CatPheo-a inhibits the telomerase *in vitro* at low micromolar concentrations being much more active than its precursor, anionic Pheo-*a*. IC_50_ values of these compounds in the modified TRAP assay [17] are 7.4 μM and > 40 μM, respectively (from unpublished results). Therefore it seems appropriate to investigate the features of CatPheo-*a* interaction with the nucleotide sequence corresponding to the human telomeric DNA, as well as to check its ability to generate singlet oxygen.

The subject of this study is CatPheo-*a* binding to Na^+^-stabilized 22-mer sequence of human telomeric DNA 5′-d[AG_3_(T_2_AG_3_)_3_]-3′ (Tel22) containing three telomeric repeats TTAGGG. It is known that in the presence of monovalent cations (K^+^, Na^+^) single-stranded Tel22 can fold into quadruplex structures of different conformations and topologies. In Na^+^-containing solutions it forms the intramolecular G-quadruplex with antiparallel basket-type conformation (PDB code 143D) where three G-tetrads are connected with one diagonal and two lateral (TTA) loops [18]. In our work the spectroscopic properties of the complexes formed between CatPheo-*a* and Tel22 were studied in neutral aqueous buffered solutions, pH6.9, of near-physiological ionic strength at various polymer/ligand concentration ratios, *P/D*, ranged from 0 to 7. An effect of the dye on the thermodynamic parameters of Tel22 folding was established, and efficiency of singlet oxygen generation by CatPheo-*a* was tested.

## 2. Materials and methods

### 2.1. Materials

The CatPheo-*a* was synthesized from Pheophorbide-*a* (Frontier Scientific Inc., Logan, Utah) according the procedure previously described in [8]. 22-mer oligonucleotide 5′-d[AG_3_(T_2_AG_3_)_3_]-3′ of human telomeric DNA (Tel22, PDB ID: 143D) from Eurogentec SA (Belgium) was used without further purification.

The 2 mM phosphate buffer (pH6.9) containing 0.5 mM EDTA and 0.1 M NaCl prepared from deionized water was used as solvent.

Quadruplex structure of Tel22 was formed by heating the solution of oligonucleotide in a buffer containing 0.1 M NaCl at 90°C for 5 min followed by cooling to room temperature in accordance with the procedure [19].

Tel22 concentration was determined spectrophotometrically using the extinction value of ε_260_ = 228500 M^−1^ cm^−1^ (in strands) at room temperature. The concentration of the dye was determined gravimetrically using CatPheo-*a* molecular weight value of 736.9 Da. It was 10 μM in all samples.

### 2.2. Apparatus and techniques

The spectroscopic properties of CatPheo-*a* and its complexes with Tel22 have been studied using polarized fluorescence and absorption spectroscopy. Effect of CatPheo-*a* on thermodynamic parameters of Tel22 quadruplex formation was determined using absorption melting techniques. The singlet oxygen generation by this dye was studied using near infra-red (NIR) spectroscopy. The sample solutions were filled in the quartz cells.

Electronic absorption spectra were recorded on SPECORD UV/VIS spectrophotometer (Carl Zeiss, Jena).

Fluorescent measurements were carried out on a laboratory spectrofluorimeter based on the DFS-12 monochromator (LOMO, Russia, 350-800 nm range, dispersion 5 Å/mm) by the method of photon counting as it was described earlier [20,21]. The fluorescence was excited by polarized beam of He-Ne laser (λ_exc_ = 633 nm) which radiation was attenuated by neutral density filters. The emission was observed at the 90° angle from the excitation beam. The fluorescence intensity, *I*, and polarization degree, *p*, have been calculated using the formulas [22]:

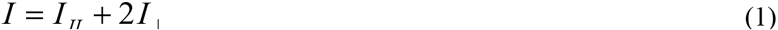

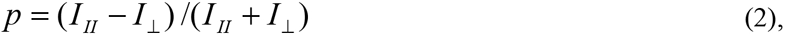

Where *I _II_* and *I*_┴_ are intensities of the emitted light, which are polarized parallel and perpendicular to the polarization direction of exciting light, respectively.

Generation of the singlet oxygen by CatPheo-*a* and its parent compound, Pheo-*a*, have been studied in ethanol solution at the same experimental conditions using laboratory setup for measurements in NIR region based on IRS-31 monochromator (LOMO, Russia) equipped with InGaAs photodiode detector. The laser diode with λ_exc_ = 665 nm (12 mW) was used for the luminescence excitation.

Binding of the CatPheo-*a* to Tel22 was studied in titration experiment where dependences of the dye fluorescence intensity and polarization degree on the molar phosphate-to-dye ratio, *P/D*, were measured under the constant dye concentration (10 μM). For this the dye sample was added with increasing amounts of the concentrated stock solution of Tel22 quadruplex containing the same dye content. Wavelength of observation was λ_obs_ = 677 nm. *P/D* values were in the range from 0 to 7.

Absorption melting experiments were carried out at 295 nm in accordance with the procedure [23]. The quartz cell was allocated in the copper cell which temperature was changed by computer operated Peltier element. The cell was inserted into the spectrophotometer. The accuracy of the temperature measurement was 0.1 °C.

All experiments were carried out at room temperature (20 – 22 °C).

## 3. Results and discussion

### 3.1. Absorption and fluorescent properties of CatPheo-a in a free state and bound to Tel22 quadruplex

Visible absorption spectra of CatPheo-*a* in a free state and bound to Na^+^-containing Tel 22 oligonucleotide measured in aqueous solution at near-physiological ionic conditions are presented in figure 2. Absorption spectrum of the free dye consists of the intense Soret band (B-band) located at 380 nm (ε_380_ = 34200 M^−1^cm^−1^) and four Q-bands [8] among which the most intensive peak is longwave one with maximum at 682 nm is (ε_682_ = 16200 M^−1^cm^−1^), that is typical for all chlorine derivatives [1,2]. These bands correspond to π→π^*^ transitions polarized within the plane of conjugated macrocycle. The porphyrin Soret band results from superposition of two mutually perpendicularly polarized intense electron transitions [24].

**Figure 2.**
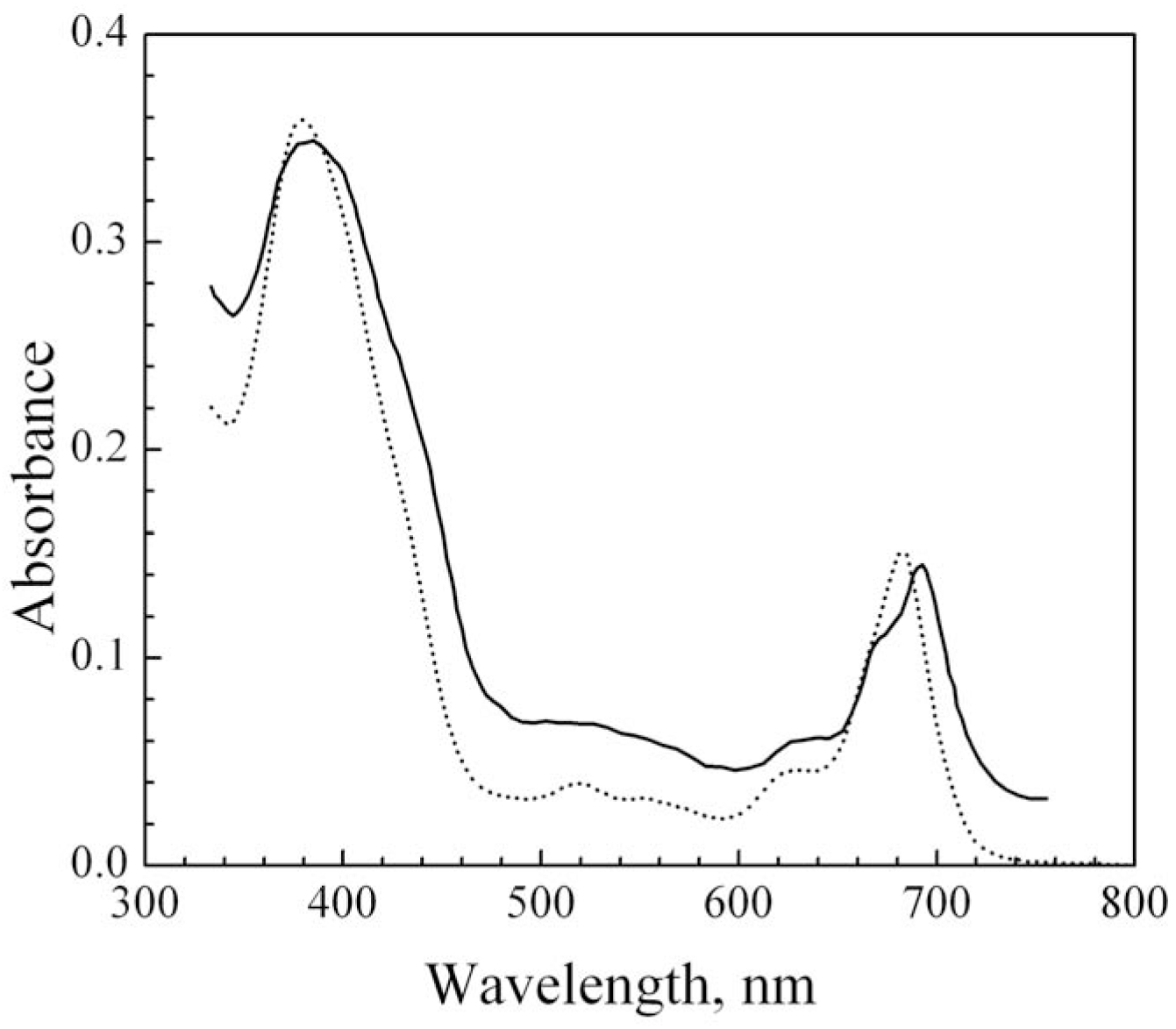
Absorption spectra of CatPheo-*a* in a free state (dotted line⋅) and bound to Tel22 quadruplex at *P/D* = 4.5 (solid line) measured in 2 mM phosphate buffer containing 0.5 mM EDTA and 0.1 M NaCl, C_dye_ = 10 μM, light path 1 cm.

The fluorescence spectrum of free CatPheo-*a* represents a broad band with a maximum at 668 nm and a longwave shoulder [8]. It can be fitted by the sum of two Gaussian components centered at 668.5 and 702.5 nm corresponding to different porphyrin states with transition dipoles polarized along X and Y direction of macrocycle [25,26]. Fluorescence polarisation degree measured in the band maximum is *p* = 0.015. Since at working concentration of 10 μM CatPheo-*a* is mainly in dimeric form [9], absorption bands can be attributed to the dye dimers, whereas emitting species is a monomer. This explains the fact that fluorescence band maximum is positioned in more shortwave region relatively to the longwave absorption band. It was shown that disintegration of the CatPheo-*a* dimers starts only at temperature more than 70 °C [9]. This process was accompanied by substantial increase in intensity of the dye absorption and emission.

Binding of CatPheo-*a* to Tel22 results in the transformation of the dye spectra. At *P/D* = 4.5 absorption of the bound dye exhibits small hypochromism, red shift and broadening of the bands in comparison with that observed for unbound one (figure 2). So the Soret absorption band undergoes 4 nm red shift (to 384 nm), whereas longwave Q-band splits into two components and shifts to 692 nm (10 nm red shift of the maximum). Appearance of the background on the edges of the spectrum evidences an increase in the solution turbidity. This phenomenon can be caused by formation of dye-oligomer aggregates at near-stoichiometric (by charge) condition that induces appearance of the light scattering. Insignificant value of hypochromicity observed in maxima of the Soret and Q-bands upon binding to Tel22 can be explained by the fact that free CatPheo-*a* (at *P/D* = 0) is in the form of dimers, which absorption is almost 2 times less than for monomers [9].

At the same time, binding to Tel22 results in substantial enhancement of CatPheo-*a* emission, rise of the fluorescence polarisation degree up to 0.2, as well as 14 nm red shift of fluorescence band maximum (to 682 nm) that can indicates withdrawal of the dye molecules from aqueous environment and formation of complex with π-stacking between their chromophores and nucleic bases.

It was shown that CatPheo-*a* is able to produce singlet oxygen in ethanol solution under irradiation by light with λ_exc_ = 665 nm. The spectrum represents a single band in near IR range with maximum at 1272 nm with full width at the half maximum amounted by approximately 20 nm. The relative quantum yield of singlet oxygen produced by CatPheo-*a* is approximately 15 % higher than that for anionic Pheo-*a* at the same experimental condition. However, no generation of singlet oxygen was detected in D_2_O solution, probably, due to dimerization of the dye.

### 3.2. Fluorescent titration study of the CatPheo-a binding to Tel22 quadruplex

Binding affinity of CatPheo-*a* to Tel22 antiparallel quadruplex was studied in fluorescence titration experiments. Figure 4 shows the dependence of the dye fluorescence intensity and polarization degree measured at λ_obs_ = 677 nm on relative concentration of Tel22 in solution within *P/D* range from 0 to 7. From the figure it is seen that the titration curve is biphasic that points out on existence of two different binding modes. In first (at *P/D* < 0.7), formation of complex between the cationic dye and the anionic oligonucleotide is accompanied by slight quenching of its emission, and then (at *P/D* > 1) it followed by the fluorescence enhancement. The fluorescence polarization degree, *p*, gradually increases reaching 0.2 at *P/D* = 7 and further continues to rise.

**Figure 3.**
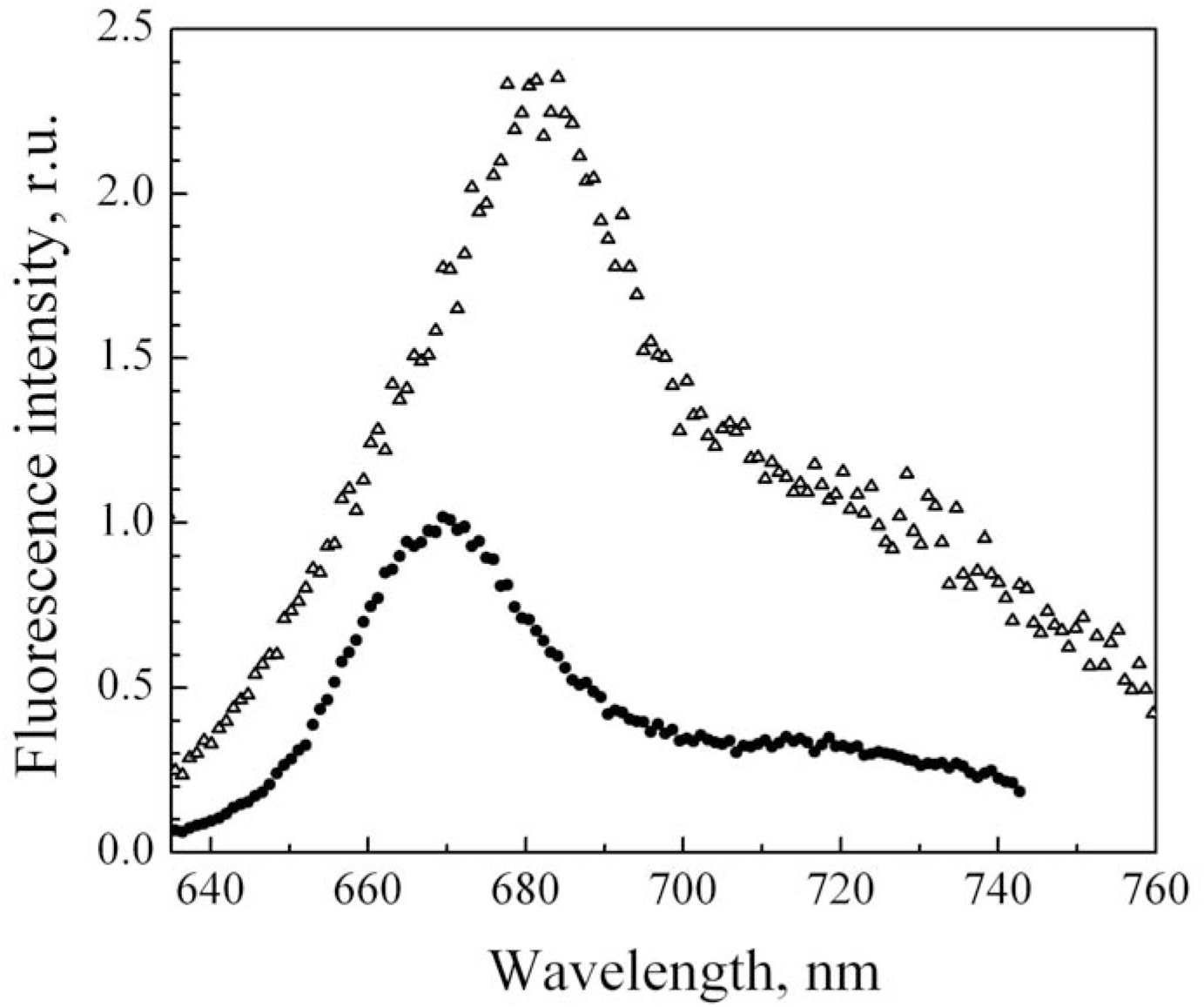
Fluorescence spectra of CatPheo-*a* in a free state (•) and bound to Tel22 quadruplex at *P/D* = 4.5 (**Δ**) in 2 mM phosphate buffer containing 0.5 mM EDTA and 0.1 M NaCl, C_dye_ = 10 μM, λ_exc_ = 633 nm.

**Figure 4.**
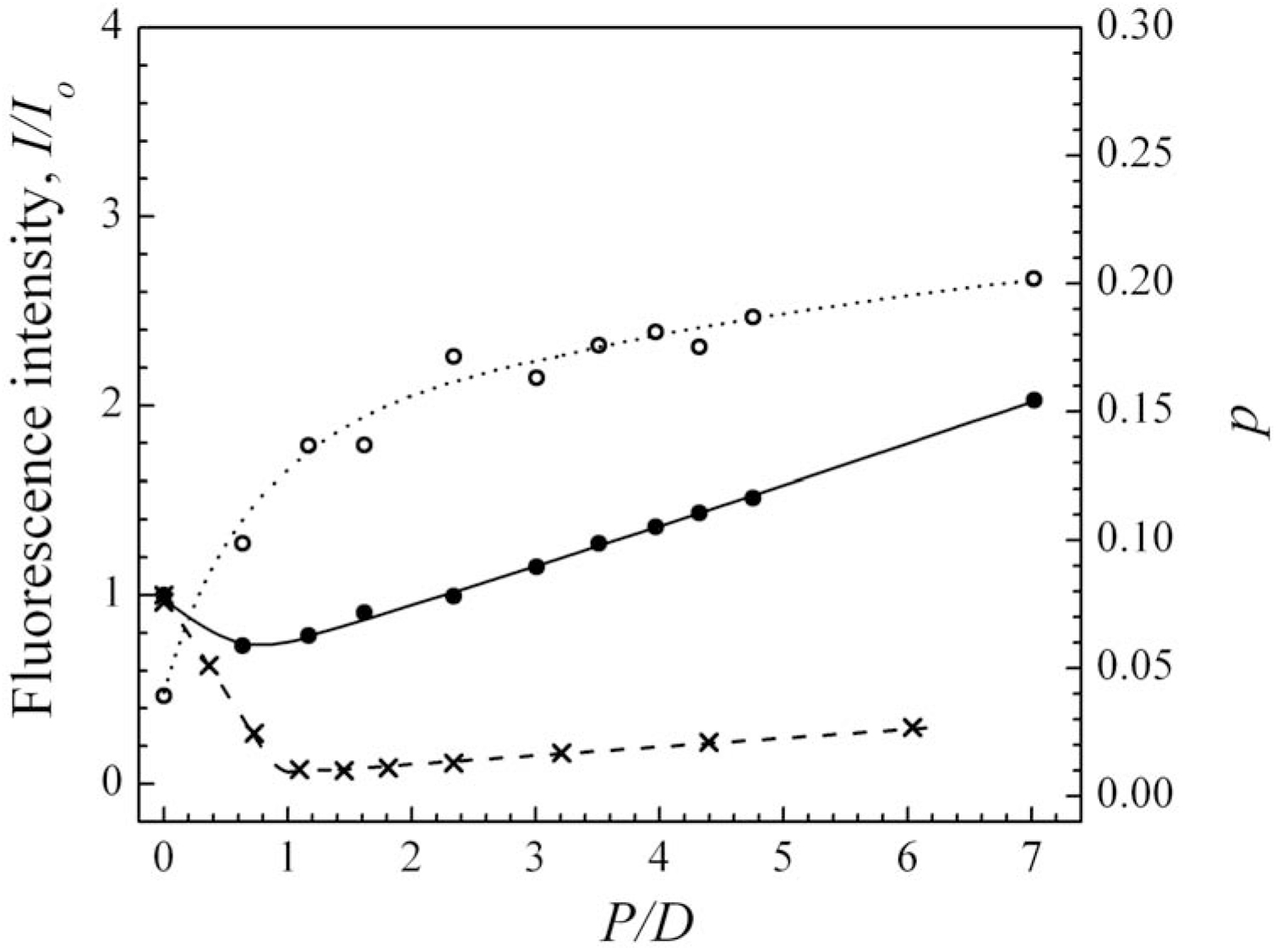
Dependence of relative fluorescence intensity, *I/I_0_*, (•) and fluorescence polarization degree, *p*, (○) of CatPheo-*a* on *P/D* molar ratio upon the titration by Tel22 complex. The measurements were carried out in 2 mM phosphate buffer containing 0.5 mM EDTA and 0.1 M NaCl, C_dye_ = 10 μM, λ_exc_ = 633 nm, λ_obs_ = 677 nm. For comparison the dependence of *I/I_0_* vs *P/D* is presented (×) for complex formed by CatPheo-*a* with poly(G) [9].

As it was reported earlier in Refs. [27,28], at low *P/D* ratios cationic porphyrins interact with linear polyanions of nucleic acids presumably by the mechanism of outside electrostatic binding to its phosphate backbone with/without self-stacking. This process is generally cooperative, and in the case of long polyanions (DNA, synthetic polynucleotides etc.) formation of the extended porphyrin stacks on their exterior usually results in the substantial quenching of the dye fluorescence [29,30]. So, for CatPheo-*a* bound to long quadruplex poly(G) strong decrease of emission intensity was observed by us at *P/D* < 1 [9]. It was characterized by linear dependence of *I/I_0_* on *P/D* and residual level of the dye fluorescence (at *P/D* = 1) of 7.5% in comparison with that for the free dye (figure 4). In the case of relatively short oligonucleotides the size of the porphyrin stacks is limited by the oligomer length and fluorescent quenching can be less. So for Tel 22 oligonucleotide which folds to form 3 guanine tetrads the stack can not contain more than 3 porphyrin molecules bound to the phosphates. Furthermore, in aqueous solution free CatPheo-*a* at *P/D* = 0 exists mostly in the dimeric form, for which the fluorescence is already partially quenched. In such a way, if we suppose that emission changes observes in the present study at *P/D* < 0.7 is conditioned by outside electrostatic binding of charged CatPheo-*a* molecules to Tel22 phosphates with self-stacking of their chromophores, then insignificant value of the fluorescence quenching observed for CatPheo-*a* + Tel22 in the initial range of titration curve (figure 4) can be explained by similar emissivity of the free dye dimers and short stacks of the dye bound to Tel22. Similar shape of the titration curve and a magnitude of the fluorescence quenching at low *P/D* ratios was observed by us earlier upon binding of porphyrin-imidazophenazine conjugate, TMPyP^3+^–ImPzn, to the same Tel22 antiparallel quadruplex [31]. In a free state this conjugate formed internal heterodimer with stacking of the chromophores and addition of the oligonucleotide solution was assumed to induce the disintegration of these dimers with subsequent binding of their porphyrin moieties to Tel 22 phosphates accompanied by macrocycle self-stacking.

The second binding type prevails at *P/D* > 1 and is characterized by the enhancement of CatPheo-*a* emission (figures 4, 3) with further increase in the dye fluorescence polarization degree. Increase in the fluorescence intensity (*I/I_0_* = 2 at *P/D* = 7) was accompanied by 14 nm red shift of emission band maximum (figure 3) that can indicate a withdrawal of the CatPheo-*a* molecules from water environment. The high value the fluorescence polarization magnitude of (*p* = 0.2 at *P/D* = 7) points out the hindering of rotation motion of bound molecules. Unfortunately, the methods used can not provide precise information one the orientation of dye molecules with respect to G4. This mode could be attributed to either the interaction of single CatPheo-*a* macrocycles with guanine tetrads through π-stacking (for example, through the end stacking with the terminal G4 tetrad) or their intercalation between the tetrads, or to incorporation of the macrocyclic chromophore into the Tel22 quadruplex groove, i.e. the type of binding remains questionable. From the literature data [32] it is known that in the antiparallel basket-type quadruplex formed by Tel22 in Na^+^–containing solution both terminal G-tetrads are hindered either by the lateral loops or by the diagonal loop that runs across the terminal G-tetrad and whose bases stack onto it. However, it may be possible that the ligand can displace the obstructing bases of the loops or bind to other elements of the G-quadruplex [32]. Perhaps, it can bind to the quadruplex grooves which in this conformation of Tel22 are accessible for dye molecules and the largest of them reaches 18.4 Å in width [18].

Substantial enhancement of porphyrin fluorescence at high *P/D* ratios was observed earlier upon binding of cationic and neutral Pheophorbide-*a* derivatives to quadruple poly(G) [9, 33], as well as for complexes formed by TMPyP^3+^ porphyrin with poly(G) [34] and Tel22 [31], porphyrin-phenazine conjugate metalized by Zn^2+^ and Mn^3+^ with poly(G) and Tel22 [35, 31]. At the same time no enhancement of CatPheo-*a* emission was observed upon its binding to double strand poly(G)⋅poly(C) and poly(A)⋅poly(U) [9], as well as to single strand poly(P) biopolymers [8]. Therefore, the fluorescence intensity was increased upon binding of these porphyrin derivatives just to G-quadruplex structures.

An existence of different types of CatPheo-*a* binding to Tel22 quadruplex is conditioned by the specific features of its structure. Planar macrocyclic chromophore of CatPheo-*a* provides π-π stacking between them and guanine quartets (end stacking with terminal tetrad, incorporation into the quadruplex groove, or, less likely, intercalation between the adjacent tetrads). The process is controlled by hydrophobic and van der Waals interactions and dominates at the polymer excess (high *P/D*). At the same time, side chain of CatPheo-*a* containing positively charged trimethylammonium group provides electrostatic interaction of the dye with negatively charged phosphate backbones of oligonucleotide which can be accompanied by formation of chromophore stacks or π-stacked complexes of macrocycle with nucleic bases in the loops. This binding type prevails at the dye excess (low *P/D*).

### 3.3. Effect of CatPheo-a on Tel22 folding: absorption melting study

To determine whether CatPheo-*a* affects the folding of Tel 22 oligonucleotide thermal denaturation of the samples were performed. Figure 5 shows UV melting curves registered at 295 nm both for Tel22 alone and for such its mixture with CatPheo-*a* (*P/D* = 0.7) where pheophorbide compound is in excess and the oligonucleotide lattice is saturated with the dye. It is seen that both of the curves are monophasic.

**Figure 5.**
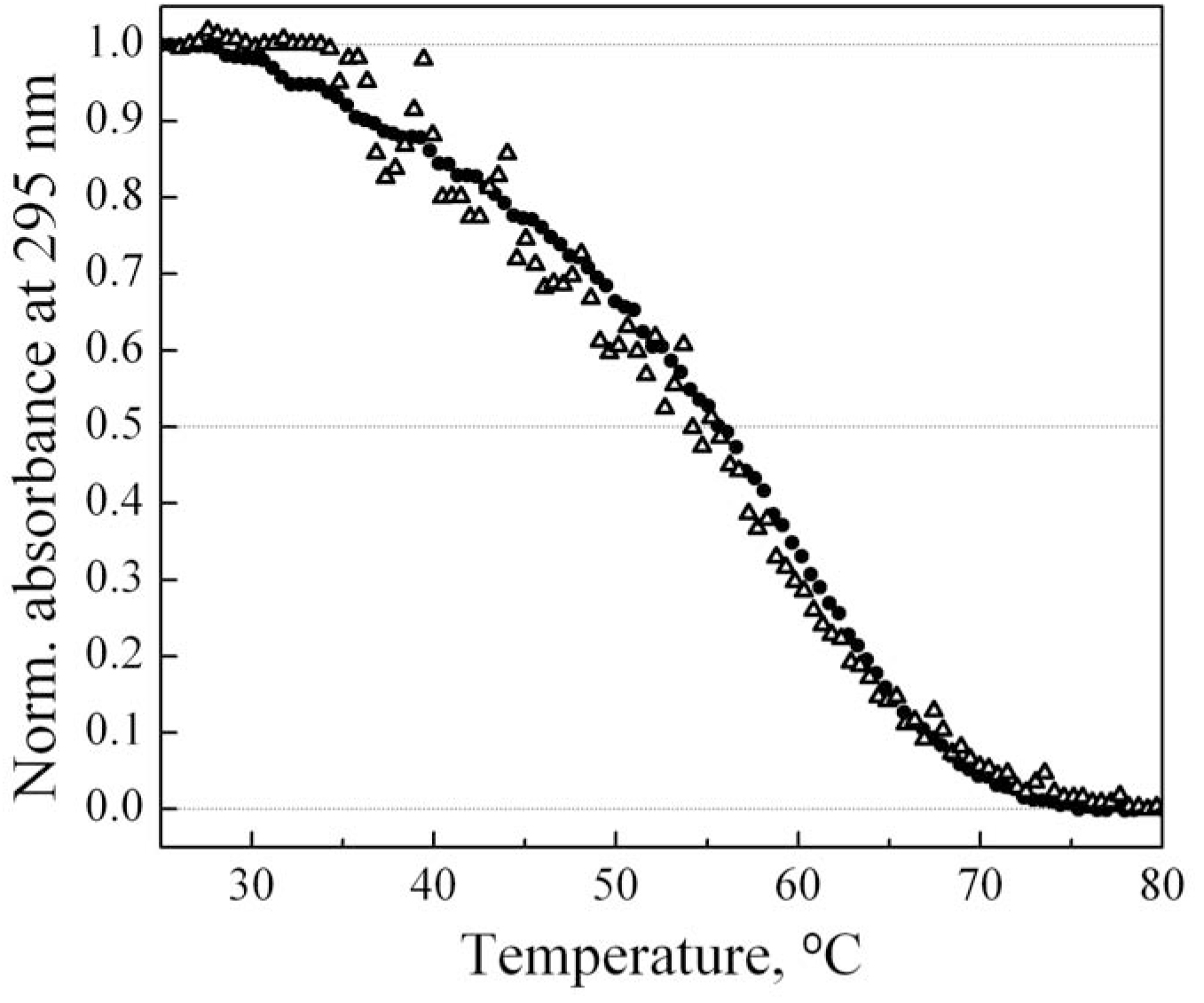
Normalized absorption melting profiles registered at 295 nm for Tel22 alone (•) and for its complex with CatPheo-*a* at *P/D* = 0.7 (**Δ**). The measurements were carried out in 2 mM phosphate buffer containing 0.5 mM EDTA and 0.1 M NaCl. Oligonucleotide concentration was 30 μM (in strands).

They have sigmoid shape and exhibit hypochromic effect at the wavelength of observation. The curves corresponding to quadruplex → single strand conformational transition (4→1) were constructed from the melting data using the standard procedure [36] as it is shown in figure 6. Here the fraction of folded oligonucleotide (i.e. fraction of the oligomer strands forming G-quadruplex), *Θ*, was calculated using the equation:

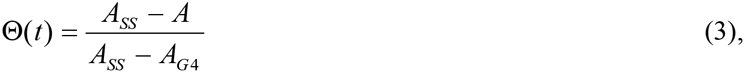

**Figure 6.**
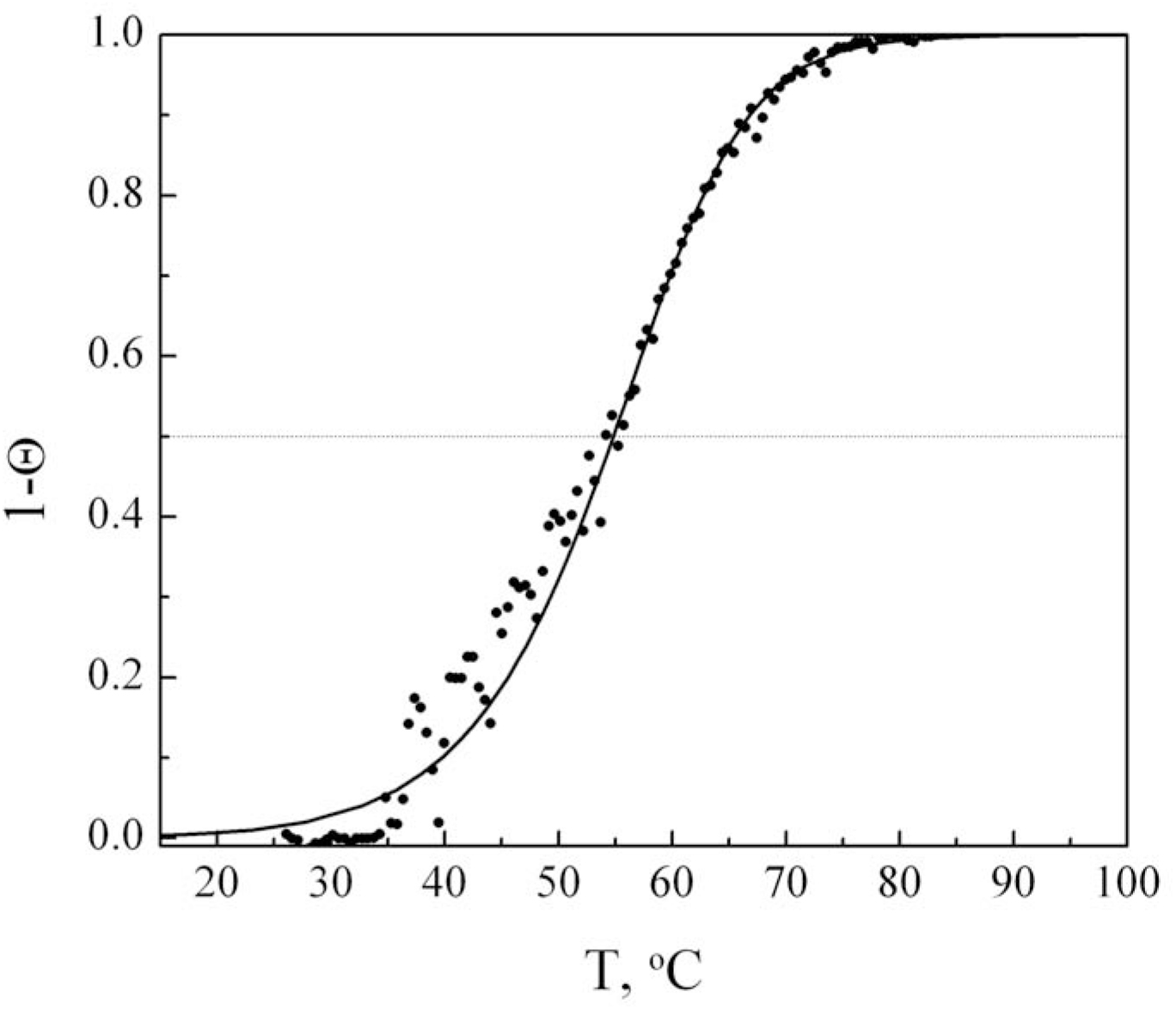
Dependence of fraction of unfolded Tel22 molecules on the temperature in the presence of CatPheo-*a (P/D* = 0.7). Solid line represents the best fits to the experimental data obtained using two-state binding model.

Where *A_SS_* – absorption of totally denatured complex, *A_G_*_4_ – absorption of the quadruplex sample, *A* – current absorption value.

To estimate changes in the thermodynamic parameters of Tel22 quadruplex folding induced by CatPheo-*a* binding the experimental transition curves corresponding to samples with and without dye were fitted to the equation based on two-state binding model [36]:

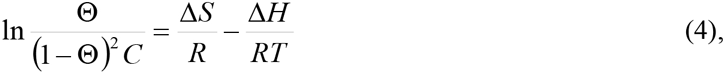

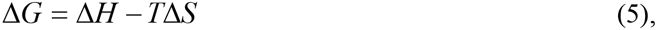

where *C* is total molar concentration of Tel22 (in strands), Δ*H* is enthalpy change, Δ*S* is entropy change, Δ*G* is Gibbs standard free energy change.

Determined thermodynamic values were summarized in the Table 1 along with transition midpoint temperatures, T_m_. As can be seen from the table, the thermally induced conformational transition from quadruplex to single strand (4→1) for individual Tel22 has T_m_ = 55.5 °C. This value is in agreement with T_m_ = 56 °C reported in [37] for the same system. It is seen, that cationic pheophorbide slightly decreases the transition midpoint temperature to T_m_ = 54.7 °C. Also the dye binding to Tel22 slightly changes the enthalpy and entropy of 4→1 transition, gives destabilizing increment in the Gibbs standard free energy calculated at physiological temperature 37 °C that results in a 2-fold reduction of apparent equilibrium constant for Tel22 quadruplex folding.

**Table 1.** Thermodynamic parameters of G-quadruplex formation by Tel22 sequence in the absence and presence of CatPheo-*a*

| Compound | $T_m$ (°C) | C (μM)<sup>a</sup> | $P/D$ | $-\Delta H$<br>(kcal/mol) | $-\Delta S$<br>(cal/mol·K) | $-\Delta G^b$<br>(kcal/mol) | $K_{eq}^c$<br>(M<sup>-1</sup>) |
| --- | --- | --- | --- | --- | --- | --- | --- |
| Tel22 | 55.6 | 17.5 | 0.0 | 52 | 134.9 | 10.15 | $1.4 \cdot 10^7$ |
| CatPheo-*a* +<br>Tel22 | 54.7 | 30.0 | 0.7 | 51.8 | 135.8 | 9.69 | $6.7 \cdot 10^6$ |

<sup>a</sup> In strands
<sup>b</sup> Data were calculated at T = 310 K using equation $\Delta G = \Delta H - T\Delta S$
<sup>c</sup> Data were calculated at T = 310 K using equation $K_{eq} = e^{-\Delta G/RT}$

From literature data it is known that some other porphyrin derivatives, in particular, tetracationic derivative TMPyP4, destabilize quadruplex structures formed by some oligonucleotide sequences [38–40]. For example, binding of the two molecules, where substituents (methyl groups) are introduced at *orto-* position of pyridyl rings, to d(TTAGGG)_4_ oligomer, which is similar to Tel22, decreases the thermostability of G-quadruplex reducing its T_m_ by 8°C [38]. Also this compound unfolds the extremely stable G-quadruplex in MT3-MMP mRNA [39] and destabilizes the tetraplex structures of d(CGG)_n_ [35]. Ni-TMPyP4 complex unfolds the quadruplex structure of Tel22 in Na^+^ containing solution [41].

Also, it is known that binding of porphyrin dyes to G-quadruplexes can induce changes in their conformation. For example, TMPyP4 promoted the conversion of Tel22 quadruplex from the basket to the hybrid type in K^+^ containing solution [42]. So, it can be one of possible reasons for G4-destabilizing effect of CatPheo-*a*.

In such a way, CatPheo-*a* efficiently binds to quadruplex DNA, however, it is not a G-quadruplex stabilizer. At the same time, its ability to efficiently generate singlet oxygen makes it a promising photosensitizer for PDT of tumors.

## 4. Conclusions

In the present study we have focused on binding of cationic Pheophorbide-*a* derivative to Na^+^-stabilized 22-mer sequence 5′-d[AG_3_(T_2_AG_3_)_3_]-3′ of human telomeric DNA (Tel22) forming an antiparallel quadruplex (PDB ID: 143D) and effect of the dye on the Tel22 quadruplex unfolding. CatPheo-*a* has much improved water solubility as compared with parent Pheo-*a* dye. The positively charged side group of this compound facilitates its binding to negatively charged nucleic acids. Investigation has been performed at various [CatPheo-*a*]/[DNA bases] ratios using several spectroscopic techniques.

It was shown that CatPheo-*a* efficiently binds to Tel22 G4 structure. Availability of two competitive binding modes was established from the biphasic character of fluorescence titration curve. The first one prevails at *P/D* < 1 and manifests itself by the fluorescence quenching. It is supposed to be outside binding of CatPheo-*a* to Tel22 phosphates accompanied by the formation of the short dye stacks on the quadruplex exterior. Not very high value of the fluorescence quenching can be explained by dimerization of the free dye. The second binding type predominates at *P/D* > 1 and is accompanied by enhancement of the dye emission as well as 14 nm red shift of the fluorescence band. Increase in the fluorescence polarization degree up to 0.2 indicates the restriction of rotation motion for the bound dye molecules. The transformations in the absorption spectrum are less pronounced (4 and 10 nm for Soret and longwave Q-band correspondingly).

The effect of CatPheo-*a* on thermodynamic parameters of Tel22 quadruplex folding was determined from absorption melting experiments. It was found that CatPheo-*a* ligand slightly destabilizes the quadruplex structure decreasing its midpoint 4→1 transition temperature by approximately 1°C. Destabilizing increment in the Gibbs standard free energy of the quadruplex folding at physiological temperature of 37°C through the dye binding was estimated as 0.46 kcal/mol that results in the 2-fold decrease of the corresponding apparent equilibrium binding constant from 1.4⋅10^7^ to 6.7⋅10^6^ M^−1^.

In ethanol solution the dye exhibits 15% higher efficiency of singlet oxygen generation as compared to the parent Pheo-*a* compound that suggests its improved photodynamic activity. So CatPheo-*a* can be considered a promising photosensitizer for PDT of tumors. These data reinforce the potential applications of pheophorbides.

The data on CatPheo-*a* binding to G4 DNA reported in this study could be helpful for rational design of novel porphyrins and structurally related compounds with enhanced affinity for telomeric quadruplex structures as potential telomerase inhibitors with antitumor properties.

## COMPLIANCE WITH ETHICAL STANDARDS

The authors declare that our manuscript complies with the all Ethical Rules applicable for this journal and that there are no conflicts of interests.

